# Dispersal: a central and independent trait in life history

**DOI:** 10.1101/065151

**Authors:** Dries Bonte, Maxime Dahirel

## Abstract

The study of trade-offs among major life history components (age at maturity, lifespan and reproduction) allowed the development of a quantitative framework to understand how environmental variation shapes patterns of biodiversity among and within species.

Because every environment is inherently spatially structured, and in most cases temporally variable, individuals need to move within and among habitats to maximize fitness. Dispersal is often assumed to be tightly integrated into life histories through genetic correlations with other vital traits. This assumption is particularly strong within the context of a fast-slow continuum of life-history variation. Such a framework is to date used to explain many aspects of population and community dynamics. Evidence for a consistent and context-independent integration of dispersal in life histories is, however, weak. We therefore advocate the explicit integration of dispersal into life history theory as a principal axis of variation influencing fitness, that is free to evolve, independently of other life history traits.

We synthesize theoretical and empirical evidence on the central role of dispersal and its evolutionary dynamics on the spatial distribution of ecological strategies and its impact on population spread, invasions and coexistence. By applying an optimality framework we show that the inclusion of dispersal as an independent dimension of life histories might substantially change our view on evolutionary trajectories in spatially structured environments.

Because changes in the spatial configuration of habitats affect the costs of movement and dispersal, adaptations to reduce these costs will increase phenotypic divergence among and within populations. We outline how this phenotypic heterogeneity is anticipated to further impact population and community dynamics.

## Introduction

Survival, fecundity and age at maturity are life-history traits that are tightly linked to population-level demography (Stearns 1989, Clutton-Brock and Sheldon 2010). They are typically organized in strategies, thus correlated and linked by both genetic and environmental covariation (Stearns 1989, Lindström 1999). Variation in these traits can be neatly integrated into an analytical framework to predict population persistence using, depending on the underlying driving variable(s), either classical matrix models or Integral Projection Models (IPMs) (Caswell 1990, Merow et al. 2014).

The quantification of life-history strategies is fundamental to advance ecology towards a predictive science enabling reliable forecasting, especially in light of global change (Selwood et al. 2015; Urban et al. 2016). The incorporation of quantitative genetics in IPM has for instance made it an indispensable tool to explore causes and consequences of eco-evolutionary dynamics in (semi-) wild populations (Coulson 2012, Smallegange and Coulson 2013, Vindenes and Langangen 2015). When detailed individual-based life-history metrics are unavailable, species-specific estimates are used to infer the fate of populations under environmental change. For instance, Salguero-Gómez et al. (2016), expanding Stearns (1983)’s approach to plants, found for 418 plant species that life strategies could be synthesised by two principle axes representing respectively the pace of life (fast-slow continuum) and a wide range of reproductive strategies. The authors subsequently used this framework to predict species demographic responses to (anthropogenic) disturbances, and advocate its use to study species persistence in changing environments. Species traits linked to the fast-slow continuum were also found to influence extinction threat status in birds and fishes (Jeschke and Strayer 2008).

Dispersal is defined as any movement potentially leading to gene flow (Ronce 2007), so movements beyond what Addicott et al. (1987) defines as the ecological neighbourhood. This definition lends itself to an individual-centred view on dispersal as a trait affected by environmental, genetic, and physical constraints (Clobert et al. 2009, Bonte et al. 2012). Dispersal is a complex, multivariate trait involving several stages — departure, transfer and settlement- that eventually determine net displacement between locations of birth and subsequent reproduction. Each of these stages is expected to be under selection so to reduce the overall costs of dispersal (Travis et al. 2012, Bonte et al. 2012). Dispersal is selected as an adaptation to avoid competition with relatives, reduce the risks of inbreeding, or spread risk in spatially and temporally varying environments or demography (Duputié & Massol 2013). The overall dispersal process may, however, be affected by other selective forces related to e.g. foraging or reproduction, rendering dispersal likely an important, but largely unintegrated trait in classic life history (dispersal as a by-product; see Van Dyck and Baguette 2005, Burgess et al. 2016).

In an ever changing world, populations will have heightened risks of extinction when dispersal is absent, whatever the survival rate and reproductive output of its constituent individuals (Berg et al. 2006). A life-history theory that includes dispersal is thus needed to understand the diversity of life forms on Earth and their dynamics in space and time. Connections between dispersal and other vital life history traits are referred to as dispersal syndromes (Ronce et al. 2012). Often, these syndromes are invoked to explain spatial dynamics of species and traits. For instance, the metapopulation/metacommunity framework incorporating dispersal-competition trade-offs (Hastings 1980; Tilman 1994, Yu and Wilson 2001, Leibold et al. 2004, Calcagno et al. 2006) has fostered the view that colonisers should have poor competitive abilities, and vice versa, to explain community dynamics. At the population-level, dispersal costs through reproduction trade-offs are often automatically assumed when developing models of dispersal evolution (Travis et al. 2012). Also at the individual level, dispersal is often considered integrated within the fast-slow continuum, with “fast” organisms spreading more, an assumption that still prevails in current iterations of the pace of life concept (Réale et al. 2010; Wolf & Weissing 2012). Organisms must become spatially dispersed as they grow, reproduce, and die, if they are to avoid density-dependence and maintain exponential growth (Holt 2009). Southwood (1977) developed in his seminal presidential address to the British Ecological Society a framework that integrated (stochastic) fitness expectations, dormancy and dispersal to understand the evolution of ecological strategies in relation to spatiotemporal dimensions of the habitat.

In this paper, we synthesise data and views on how dispersal is integrated into life-history strategies. We show that evidence for a consistent genetic correlation with other traits is -not unsurprisingly- weak. We plead to put dispersal independently central in life histories and discuss implications for our understanding of spatial eco-evolutionary dynamics.

## Is dispersal genetically integrated in life histories?

Dispersal syndromes appear to be present at the species-level. Their strength and direction are, however, highly inconsistent because of multiple and highly variable selection pressures involved in fitness maximisation. Across terrestrial animal species, dispersal is typically integrated into life histories through size dependency (Stearns 1983, Gaillard et al. 1989), but the relationship between dispersal and size is definitively not general, nor uniform among several taxa (Stevens et al. 2014. Most importantly, stronger evidence for dispersal-size correlations was found in aerial dispersers and ectotherms, relative to ground dwellers and endotherms. This suggests that dispersal becomes more integrated in species life histories in organisms that have specific adaptations for longer distance movements (i.e., wings). Such investments are often very costly from a developmental perspective and therefore trade-off with investments in reproduction or early-maturation. In butterflies, life history traits related to reproduction and size are good predictors of dispersal distance, but whether correlations results from genetic integration or from independent responses to environmental variability remains unknown (Stevens et al. 2013, 2014). For plants, Thompson et al. (2002) did not find convincing evidence for trade-offs between seed dispersal and several demographic parameters. Seed mass and especially plant height did contribute most to dispersal distance (Thomson et al. 2011). Seed morphology is, however, a poor predictor of (long-distance) dispersal because it is determined by multiple processes that move seeds, for instance multiple vectors (Higgins et al. 2003). In many marine organisms, dispersal is typically assumed to be associated with traits like offspring number and larval growth. Much of the variation in dispersal kernels is, however, maintained as a side product of the evolution of other vital traits, for instance for utilising pelagic habitat to acquire resources prior to metamorphosis to the sedentary life stages (Burgess et al., 2016).

Consistent genetic dispersal syndromes at the individual-level are neither expected to maximise fitness. Like life-history traits related to reproduction and aging (Lindström 1999), dispersal is now acknowledged to be context- and density dependent (Clobert et al. 2009, De Meester and Bonte 2010, Bitume et al. 2013, 2014) and thus highly heterogeneous when studied at the phenotypic level. An analysis of phenotypic data on dispersal and movement of European butterflies revealed at least as much variation within as among species (Stevens et al. 2010). At the individual level, and under environmental conditions favouring a single ESS, dispersal should always be optimised to maximise fitness under the prevailing environmental conditions (Bonte et al. 2014) although constraints due to other selection pressures might shape evolutionary trajectories in a non-adaptive manner (Burgess et al. 2015). While numerous cases of within-species life history-dispersal syndromes have been reported, there is no consistency in the strength and sign of specific associations (Ronce et al. 2012). So far, few studies have gone beyond simple observations of phenotypic correlations and evaluated the genetic integration of dispersal in individual life histories. Guerra (2011) found by meta-analysis that the strength and direction of the association between wing dimorphism and life traits was variable among species. Only relationships with fecundity (flight-oogenesis syndrome) appeared to be overall strong, although some insect orders (e.g. Coleoptera) consistently present the opposite of the “canonical” trade-off pattern. Genetic correlations between wing morph expression and fecundity have been consistently found in several wing dimorphic cricket species (Roff 1994, Roff and Bradford 1996, Roff and Fairbairn 2011).

In species lacking obvious dispersal polymorphism, dispersal syndrome evolution tends to be highly context-dependent. In the moth *Epiphyas postvittana*, the strength of genetic correlations between flight ability and lifespan/fecundity was highly variable between populations (Gu and Danthanarayana 1992). In the spider mite *Tetranychus urticae*, no genetic correlations were observed between aerial dispersal and fecundity (Li and Margolies 1994). In *Erigone* spiders and *Melitaea* butterflies, phenotypic correlations with vital rates were observed across different experimental or metapopulation contexts, pointing at the existence of colonization syndromes (Bonte and Saastamoinen 2012). In both species however, consistent correlations could not be detected. Figure 1 (data from Bonte et al. 2008) visualises the correlation of several dispersal proxies (in blue) with vital traits like reproductive output, reproduction, age at maturity and web building. Spiders were reared using a half-sib design at four different temperatures. Some dispersal proxies (activity and ballooning) correlated with age at maturity at the individual level when all treatments were combined (i.e., the arrows of Act, Bfrq, and TTM point in the same direction in the first 2 Principle Component Axes). This correlation does, however, disappear when individual traits are studied within each temperature treatment (Fig 2), i.e., the dispersal traits no longer point in the same direction as the other life history traits. In *Melitaea*, the overall phenotypic correlations equally disappeared when mobility was tested under common garden conditions (Duplouy et al. 2013).

**Fig 1.**
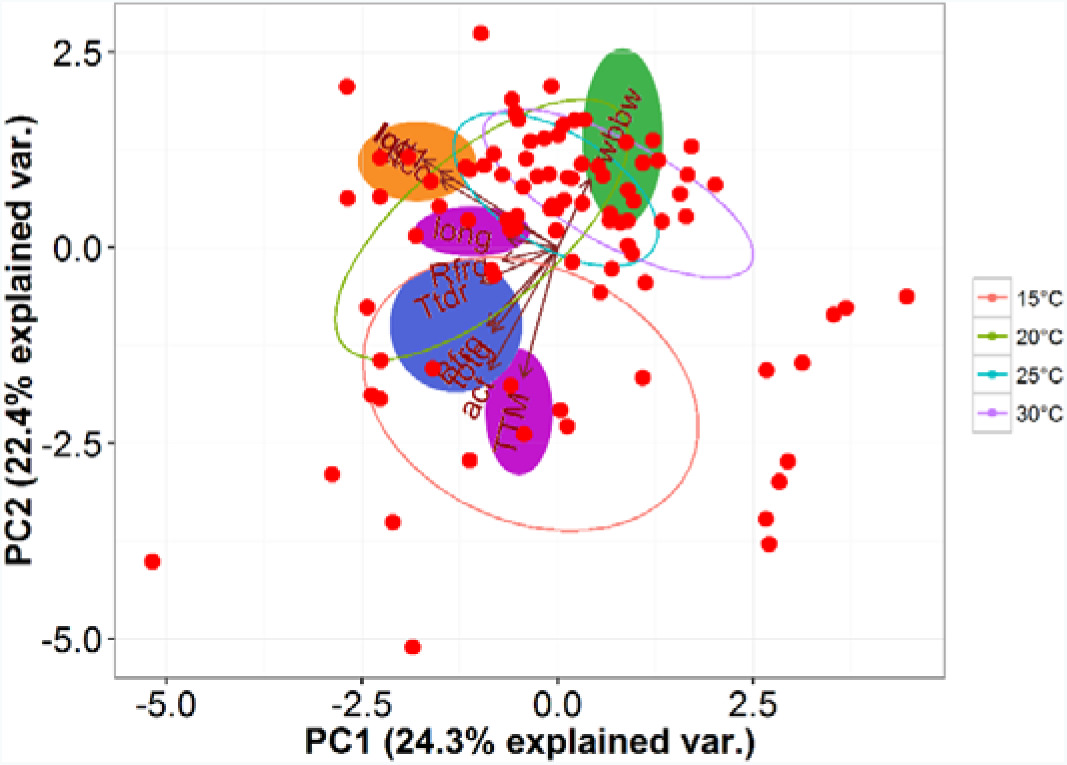
*Organisation of spider life histories along principal components. The following traits were tested for sibs of*Erigone atra *reared at four different temperatures during development conditions under laboratory conditions at the individual level: trait linked to dispersal (Ballooning frequency Bfrq, Rapelling frequency Rfrq, total duration of the tiptoe behaviour Ttdr, overall activity Act; in blue) time till maturity TTM or longevity LONG (in purple), web building behaviour (green) and fecundity (total egg sacs ttcc, lifetime fecundity lgtt and eggs/eggsacs log1; in orange). Strategies are determined by the breeding temperature conditions (data from Bonte et al. 2008)*

**Fig 2.**
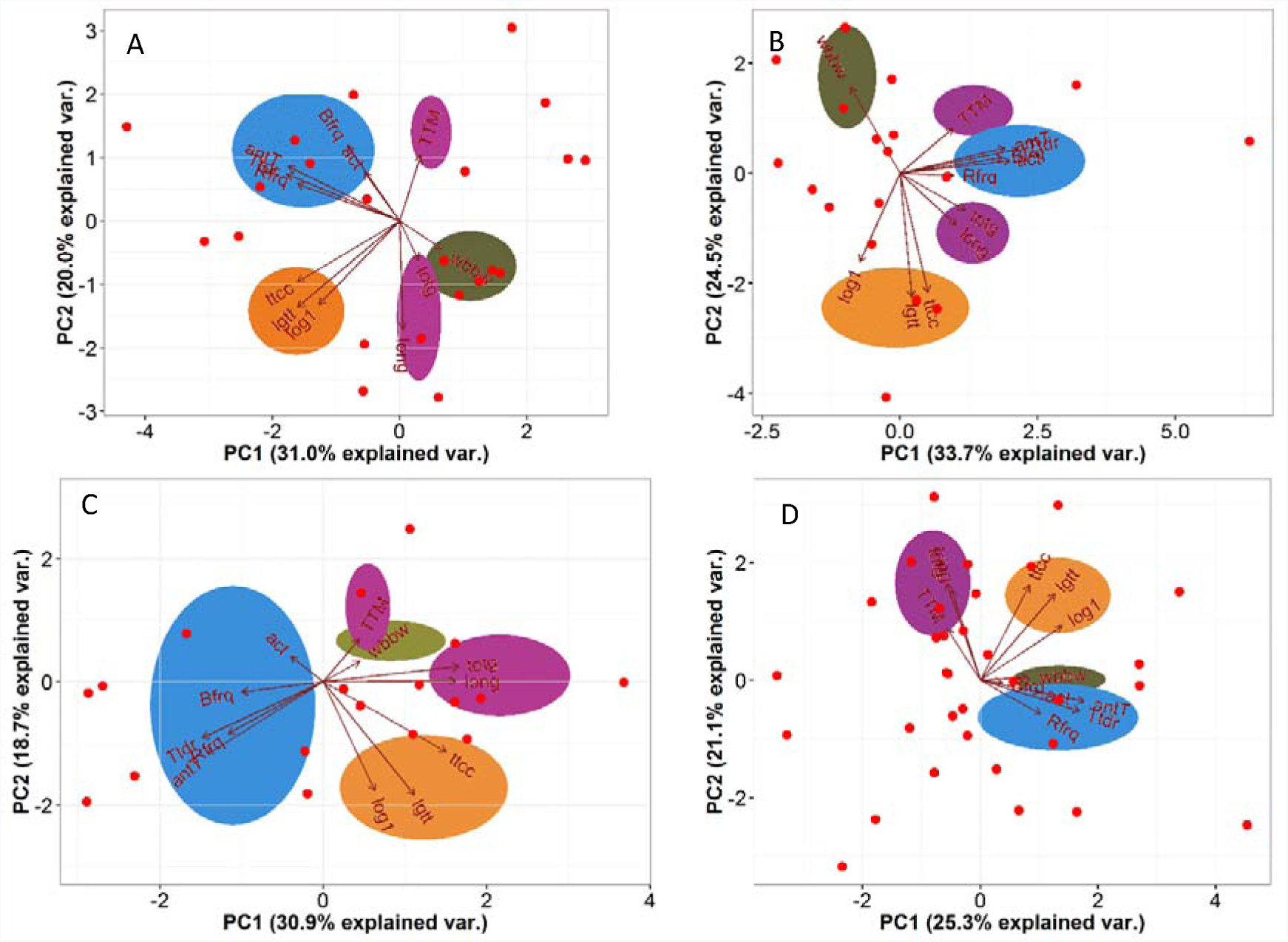
*Traits related to dispersal are differently correlated with the overall life history contingent to the temperature of development (A: 15°C; B: 20°C; C:25°C; D: 30°C). The following traits were tested for female sibs* Erigone atra *reared at four different temperatures during development conditions under laboratory conditions at the individual level: trait linked to dispersal (Ballooning frequency Bfrq, Rapelling frequency Rfrq, total duration of the tiptoe behaviour Ttdr, overall activity Act; in blue) time till maturity TTM or longevity LONG (in purple), webbuilding behaviour (green) and fecundity (total eggsacs ttcc, lifetime fecundity lgtt and eggs/eggsacs log1; in orange). Strategies are determined by the breeding temperature conditions (data from Bonte et al. 2008)*

Although there is definitely a need to collect more data on the shape, strength and consistency of individual dispersal syndromes, available theory on the joint evolution of dispersal and other traits largely confirm the optimality of independently evolving dispersal-life traits. Dytham and Travis (2006) allowed age-at-death to evolve jointly with emigration and found the cost of dispersal to strongly affect the joint and non-consistent evolution of both traits. Similarly, landscape dynamics, costs of dispersal, the level of environmental stochasticity affect both the strength and direction of dispersal-fecundity syndromes (Ronce et al. 2000, Bonte and de la Peña 2009). A special case is the study on the joint evolution of dispersal and selfing ability in plants, where the probability of finding a patch with pollinators and the cost of dispersal affected the ESS for both traits, but not necessarily in an “optimal” way (Cheptou & Massol 2009, Massol & Cheptou 2011).

Evidently, when dispersal and other life traits are genetically linked, the shape of the trade-off function will affect the mutual evolutionary dynamics (Berdahl et al. 2015, Weigang and Kisdi 2015)

## Consequences – genetic (co-)variation

The available data and theoretical insights so far point to dispersal being a central and independent life history trait, with impacts on both local and regional demography. While genetic correlations with other traits are far from general, there is compelling evidence for genetic variation in dispersal (Zera and Brisson 2012). It is therefore central to eco-evolutionary dynamics in metapopulations. Under the assumption that dispersal is only determined by genetic variation, any spatiotemporal variance in fitness expectations among patches selects for dispersal till the level where an ideal free distribution is reached (Hastings 1983, Poethke & Hovestadt 2002, Massol et al. 2011). Under such conditions, the evolution of dispersal will be relaxed until kin competition, inbreeding and demographic stochasticity become too strong. The balance between demographic variation, inbreeding and kin competition will thus maintain spatiotemporal variation in dispersal at the metapopulation-level. The close interaction between ecology and evolution in metapopulations has for instance been demonstrated in Glanville fritillary butterflies in which dispersal and overall activity is determined by allelic variance in *PGI* (a key-enzyme in metabolism) among young and old islands populations (Hanski and Saccheri 2006, Klemme and Hanski 2009, Niitepõld et al. 2009, Niitepõld 2010, Hanski and Mononen 2011).

Dispersal and traits related to growth can be selected during range expansion and eventually affect species range limits (Phillips et al. 2010, Therry et al. 2014a, b, Kubisch et al. 2014, Fronhofer and Altermatt 2015, Huang et al. 2015). This process of spatial sorting is expected to theoretically accelerate range expansion and invasions (Bénichou et al. 2012, Shine et al. 2011). The strength and direction of the genetic correlation between dispersal and other traits will, however, affect ecological and evolutionary processes in a different way (Burton et al. 2010). More-importantly, dispersal may again evolve as a side-product of, or contingent on, the evolution of other vital traits. Individual-based models show for instance that dispersal evolution during range expansion can be constrained by investments in competitive abilities, reproduction (Burton et al. 2010) and thermal adaptations (Hillaert et al. 2015). In absence of genetic correlations, dispersal and life history traits independently evolve according to respectively spatial sorting and local environmental conditions (Van Petegem et al. 2016). It remains thus to be studied to which degree evolved changes at range borders effectively affect ecological dynamics since environmental and demographic conditions may be highly different.

The absence of consistent dispersal syndromes at the species-level forces us to critically question the fundamental role of dispersal limitation as a determinant of trait and functional diversity dynamics during succession (Lortie et al. 2004). The decoupling of dispersal and vital rates at the species-level implies that colonisation dynamics will be more determined by propagule load -the regional abundance of certain species-and local filtering effects rather than by dispersal constraints *per se.* This view forces us subsequently to move away from a deterministic theory on community assembly, but to consider selection, stochasticity and contingency as the most important driving mechanisms (Vellend 2010, Vellend et al. 2014a, b). In such cases, spatial isolation rather than species-averaged dispersal traits will be a strong predictor of both local and regional trait variation. While experimental seed addition experiments (e.g., Pinto et al. 2014) highlighted the potential structuring role of dispersal limitation, inverse approaches linking trait diversity to the local environment, isolation and dispersal are anticipated to challenge current views on assembly rules in spatially structured systems that are in disequilibrium. A recent study on functional variation in island plant communities for instance did not find a strong support for the role of dispersal in explaining trait variation associated with spatial isolation, despite a close association between functional traits and dispersal mode (Negoita et al. 2015). Obviously, firm inference of mechanisms driving community assembly is difficult to draw from pattern-oriented approaches. Using simulations of expanding finite populations, Williams et al. (2016) found that expected eco-evolutionary dynamics along expanding fronts may be highly dependent on the patchiness of landscapes and that the long-term outcome of evolution is strongly depended on which strategy initially prevailed through a spatial priority effect. In the same vein, serial founder events and drift on the expanding range edge render evolutionary trajectories in the invasion vanguard highly stochastic and eventual ecological dynamics highly unpredictable (Phillips 2015). We thus advocate researchers to carefully select replicated systems that allow disentangling the respective role of determinism versus stochasticity and contingency as driving factors of community assembly. Given the link between dispersal, species communities and eventual ecosystem fluxes (see contribution Massol et al. 2017 in this issue), a similar perspective for meta-ecosystem function is warranted.

## Consequences – condition dependency

When drift and founder effects can render spatial processes highly stochastic, then to which extent can we generalise insights from the few studies that found significant genetic covariation between dispersal and life traits, or from those reporting selection during range expansion? Indeed, even established dispersal syndromes have been shown to break down or reverse when tested in a different context (Guerra 2011, Cote et al. 2013). Similarly, while evidence on the genetic background of dispersal is strong, an equal share of work has demonstrated an overruling importance of plasticity and context-dependency in many cases (Clobert et al. 2009). Generally, emigration and immigration strategies need to be considered as threshold responses towards the local environment (context) or trait value (condition) (Clobert et al. 2009). The threshold value may be heritable and depicts the developmentally or environmentally determined condition at which individuals decide to switch behavioural states (Bonte and Lens 2007, Yan et al. 2014), in this case to leave or to settle (Fig. 3). Theory predicts that different conditional dispersal strategies might evolve in response to metapopulation conditions (Bonte and de la Peña 2009, Gyllenberg et al. 2011, Kisdi et al. 2013). Based on similar optimality arguments, different phenotypes can be expected to be associated with dispersal when local environmental conditions are different, for instance in terms of density and relatedness. In addition, transgenerational plasticity (Bitume et al. 2014, Van Petegem et al. 2015), disease and more cryptic microbial interactions (Goodacre et al. 2009, Debeffe et al. 2014) may affect dispersal and its association with other vital traits. These insights further strengthen the case for leaving a deterministic view of spatial ecology, to embrace context-dependency. Such a view, however, should not imply that dispersal should be abandoned when studying mechanisms and patterns of spatial dynamics, but that its complexity should be better accounted for.

**Fig 3.**
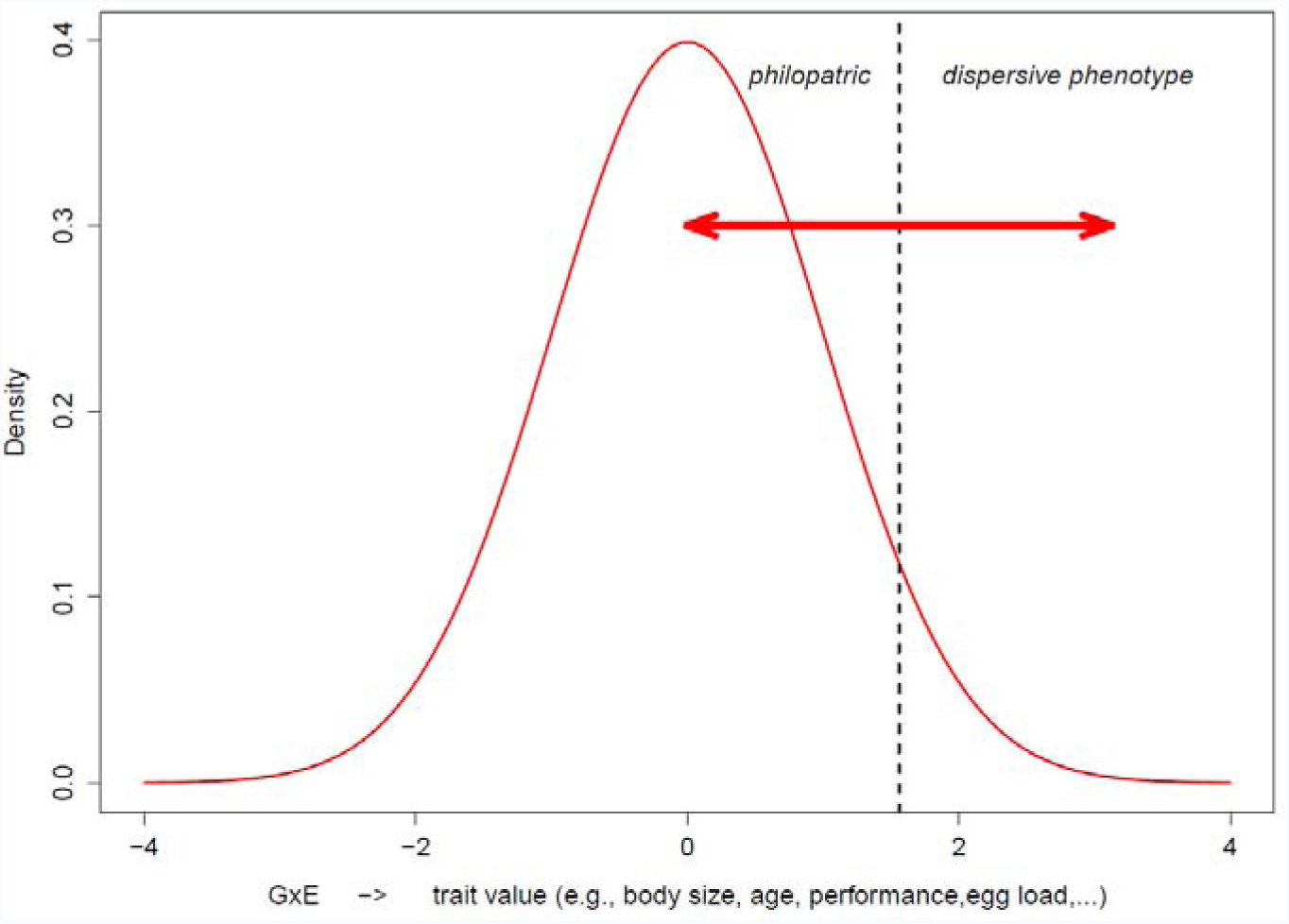
*The trait threshold model for dispersal. A population exhibits a substantial amount of phenotypic trait variation, here explanatory Gaussian distributed. Depending on the environmental conditions, density for instance, the threshold values (dashed line) for the phenotypic trait (determined by GxE) to switch between staying and leaving may shift to the right or left (red double arrow). The initial distribution is subsequently split into a divergent trait distribution of the philopatric and dispersive phenotype.*

The phenotypic signature of dispersing individuals can strongly affect metapopulation and metacommunity processes at ecological time scales (Shima et al. 2015, Jacob et al. 2015, Laroche et al. 2016). Individual phenotypic variation in size contributed for instance more to population dynamics than environmental fluctuations in several mammal species (Ezard et al. 2009). Phenotypic correlates with body size were even demonstrated to affect population dynamics more strongly in receiver populations (Ozgul et al. 2010). These advantages can be directly related to enhanced competitive abilities to take-over various resources (e.g., Bonte et al. 2011) or abilities to deal with general stressors. Furthermore, released competition at founder patches relaxes density-dependence, and may subsequently lead to monopolisation processes that control population sizes, community assembly and evolutionary potential (Urban and De Meester 2009, De Meester et al. 2016, Williams et al. 2016). Within a food web perspective, phenotypic variation has been demonstrated to affect interaction strengths between prey and predators (terHorst et al. 2010). Carry-over effects across life stages (Van Allen and Bhavsar 2014) or even generations (Bitume et al. 2011, 2014) can be particularly relevant to maintain and generate spatiotemporal variation in demography among local populations (Benard and McCauley 2008).

Phenotypic trait variation among dispersers may thus largely affect population and community dynamics (Miner et al. 2005). Just like stochastic switching strategies are advantageous in locally fluctuating environments (Acar et al. 2008), adaptive dispersal phenotype switching should lead to metapopulation stabilisation. Conditional strategies are then additionally expected to lead to maintenance of genetic and phenotypic variation within metapopulations. Both genotypic variation and phenotypic dissimilarity need therefore to be considered collectively to better understand population performance (Ellers et al. 2011). Flexible floating strategies, in which non-dispersive individuals contribute to parental fitness through helping, are another demonstration of how phenotype-dependent dispersal strategies eventually have an ecological impact beyond the individual (Cockburn 1998). Unsurprisingly, available evidence on the interplay between phenotype-dependent dispersal and demography comes from systems that allow manipulation of spatial structure, population dynamics and dispersal phenotype. By using a marine bryozoan as a model, Burgess and Marshall (2011) demonstrated interactive effects of colonizer phenotype and abundance on the reproductive yield of populations. In their study, populations established from a few individuals with short dispersal durations, which means better condition and potentially shorter distances, had similar reproductive yield as populations established by much higher number of founders that experienced long dispersal duration. In contrast, population biomass and recruitment were much influenced by colonizer abundance, but not colonizer phenotype. Context-dependent changes in disperser abundance and phenotype can thus lead to complex and poorly understood decoupling of demographic parameters. Svanfeldt et al. (2017) additionally demonstrate that changes in dispersal alter selection regimes on other traits after settlement. Finally, while habitat matching strategies may maximise metapopulation performance and connectivity (Jacob et al. 2015), converse consequences are expected in environments where selection pressures vary over spatial scales such that phenotype- environment mismatches emerge. Such habitat mismatches can be expected in marine but also many terrestrial organisms and will eventually constrain connectivity (Marshall et al. 2010)

## Conclusion

Fitness is maximised by the optimisation of life histories, usually defined in terms of survival and reproduction. Without dispersal though, any deme is doomed to go extinct. To date, dispersal is typically considered to be genetically correlated with other life history traits like reproduction or survival. We here advocate that this assumption may hold at the species level, but that dispersal needs to be considered as an independent axis in the multivariate life-history trait space at the individual level, with observed trait associations being largely context-and scale-dependent. We lack a general theory on how syndromes between dispersal and life-history traits change according to local and metapopulation-level environmental conditions. This decoupling forces us to rethink the deterministic nature of spatial population and community processes. There is evidence that phenotypic variation in dispersal can alter ecological dynamics, but more experimental work is needed to deepen insights on the role of conditional dispersal for eco-evolutionary dynamics in space.

## Acknowledgements

MD is a post-doctoral fellow funded by the Fyssen Foundation. DB acknowledges FWO research network EVENET and Belspo IAP ‘SPEEDY’ for funding and is grateful to the Nordic Oikos Society for the support to organise the thematic symposium. Scott Burgess, François Massol and Denis Réale provided very valuable comments on an earlier draft version.

